# Mining endophytic microbiome information from plant and animal transcriptome data

**DOI:** 10.1101/2021.05.07.443205

**Authors:** Guomin Han, Xianjin Wang, Guiping Qiu

**Author notes:** Author for correspondence: G. H.

## Abstract

Endophytic microorganisms play important physiological functions in plants and animals. In this paper, we developed a method to obtain endophytic microbiome information directly by analyzing transcriptome sequencing data files of plants and animals. Compared with the use of amplicon analysis or whole-genome sequencing of animal and plant tissues to analyze microbial composition information, this method can obtain endophytic microbiome information in addition to obtaining gene expression information of host plants and animals.

## Introduction

Microorganisms are almost ubiquitous, such as rivers, lakes, seas, soils, air, body surfaces, and insides of humans, animals, and plants. Many microorganisms, including bacteria, archaea, and fungi, can live in the tissues of plants. Endophytes are microorganisms that are present in various tissues of the host plant in a symbiotic or beneficial manner or without causing any detrimental effects (Ali, et al., 2021; Kaul, et al., 2016). A few endophytes can enhance the host plant tolerance to environmental abiotic stresses, such as thermotolerance (Marquez, et al., 2007; Shekhawat, et al., 2021), while others can promote plant growth through the production of phytohormones, solubilization of phosphorus and potassium, biological fixing of nitrogen, inhibition of ethylene biosynthesis (Fadiji, et al., 2021; Lata, et al., 2019). In addition, many endophytes can protect plants from microbial pathogens by the production of ammonia, hydrogen cyanide, and siderophores (Ali, et al., 2021; Carrion, et al., 2019; Lata, et al., 2019). Pathogenic fungi and symbiotic fungi can also penetrate plant cells for absorption or exchange of nutrients (Han, et al., 2019; Han, et al., 2020).

Current endophytic microorganisms detection methods mainly including culture-dependent and culture-independent methods (Dissanayake, et al., 2018; Tian, et al., 2019). Several culture-independent methods, such as Real-time PCR, In situ hybridization FISH, microarray, metabarcoding approaches, whole plant genome DNA sequencing have been used (Aslam, et al., 2017; Dissanayake, et al., 2018; Fadiji, et al., 2021; Fadiji, et al., 2021), and the mystery of a large number of plant endophytes was revealed (Aslam, et al., 2017; Matsumoto, et al., 2021; Trivedi, et al., 2020). The use of whole plant tissues for DNA extraction for amplifying 16S rDNA, fungal ITS area, or amplifying specific target genes and then sequencing with the second or third-generation sequencing technology is the often-used method. Another next-generation sequencing method is the direct sequencing of the whole genomic DNA from plant tissues and analysis using microbiome analysis software to generate endophytic microbiome information.

In this paper, a new method for mining endophytic microbiome information from plant transcriptome data was described. Briefly, after the removal of adaptor and low-quality bases, the clean data were mapped to the host plant genome or all cDNA files to generate unaligned files. The unaligned files were further analyzed by using a microbiome analysis pipeline to obtain plant endophytic microbiome information.

## Materials and Methods

### Endophytic microbiome mining method

The Workflow diagram is shown in Figure 1. The animal or plant transcriptome raw data were assessed via fastqc, and then cleaned via trimmomatic with the default parameter. The clean data were mapped to the host genome by using hisat2 with the default parameter. Use its output unaligned sequence parameter “--un-conc” to obtain an unaligned data file, which contains not only a part of the host animal and plant sequences but also expressed microbial gene fragments. Use microbiome analysis piplines, such as Kraken2, bracken and Pavian (Breitwieser and Salzberg, 2020; Lu, et al., 2017; Wood, et al., 2019), to analyze the unaligned data files to generate endophytic microbiome information.

**Figure 1.**
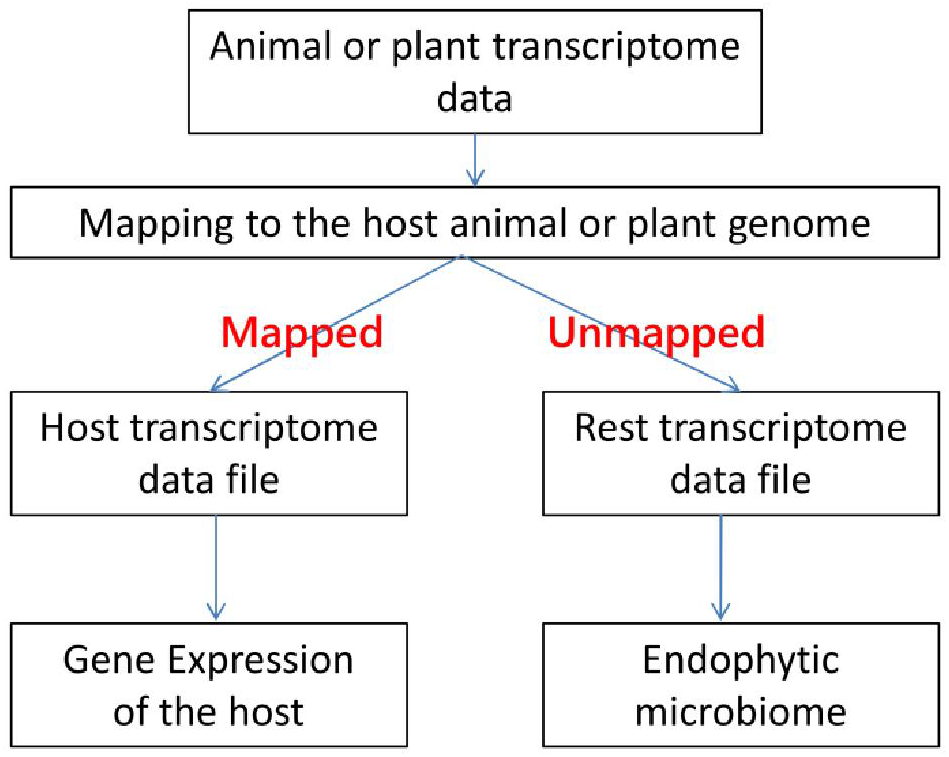
Workflow diagram

### Data sets

We selected the RNA-seq data of two plants inoculated or infected by microorganisms.

#### (1) Maize inoculated with arbuscular mycorrhiza (AM) fungus data sets

In our previous study, the maize (*Zea mays*) roots inoculated with AM fungus *Rhizophagus irregularis* DAOM-197198 (previously known as *Glomus intraradices*) were compared with the control without fungal inoculation (Han, et al., 2019). The maize raw RNA-Seq data (Bioproject accession: PRJNA553580) were used in this study. The maize inbred line B73 genome was download from Ensembl Plants.

#### (2) Common bean infected by Xanthomonas data sets

In the study by Foucher et al. (2020), a common bean (*Phaseolus vulgaris* L.) susceptible genotype (JaloEEP558) was infected by *Xanthomonas phaseoli* pv. *phaseoli* strain CFBP6546R 48 h after inoculation (Foucher, et al., 2020). The raw RNA-Seq data of Common bean (SRA accession: SRP273448) were used. The Common bean genome was download from Ensembl Plants.

## Results and discussion

### Generation the unaligned files from the host RNA-seq data

Approximately 18-28% of maize RNA-seq data cannot be mapping to the maize genome (Table S1), while about 5-11% of common bean RNA-seq data cannot mapping to the common bean genome (Table S2). The unmapping rates from maize inoculated with AM fungus are higher than that of the control, similar results were also observed in the common bean RNA-seq data (Figure 2).

**Figure 2.**
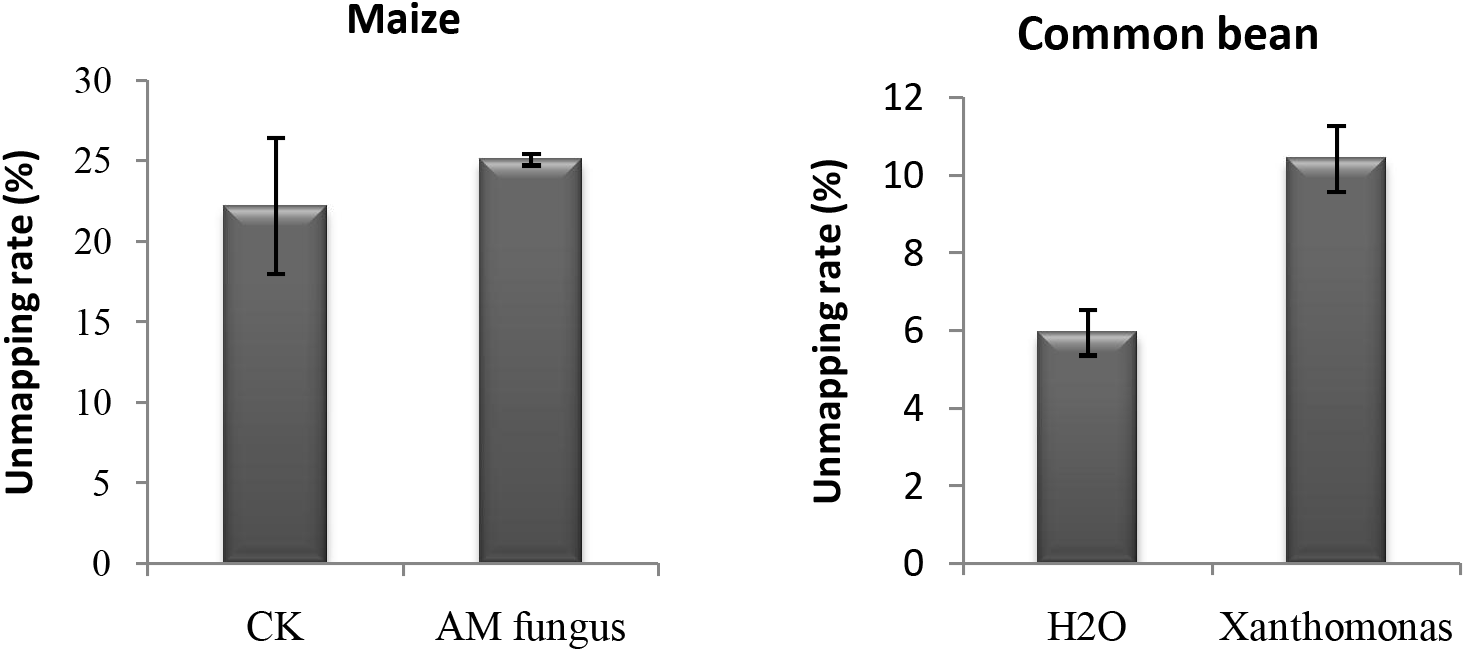
The unmapping rate of RNA-Seq data to each host genome.

### Endophytic microbiome information

As an example, in addition to maize root transcriptome data, the transcriptome data also included fungi, bacteria, archaea, and viruses, and similar results were found in the common bean transcriptome data (Figure 3). Further analysis can be done to obtain the content relationships between various types of microorganisms (Figure 4). The microorganisms in the maize root inoculated with AM fungi were mostly fungi, whereas microorganisms in common beans were dominated by bacteria (Figure 4).

**Figure 3.**
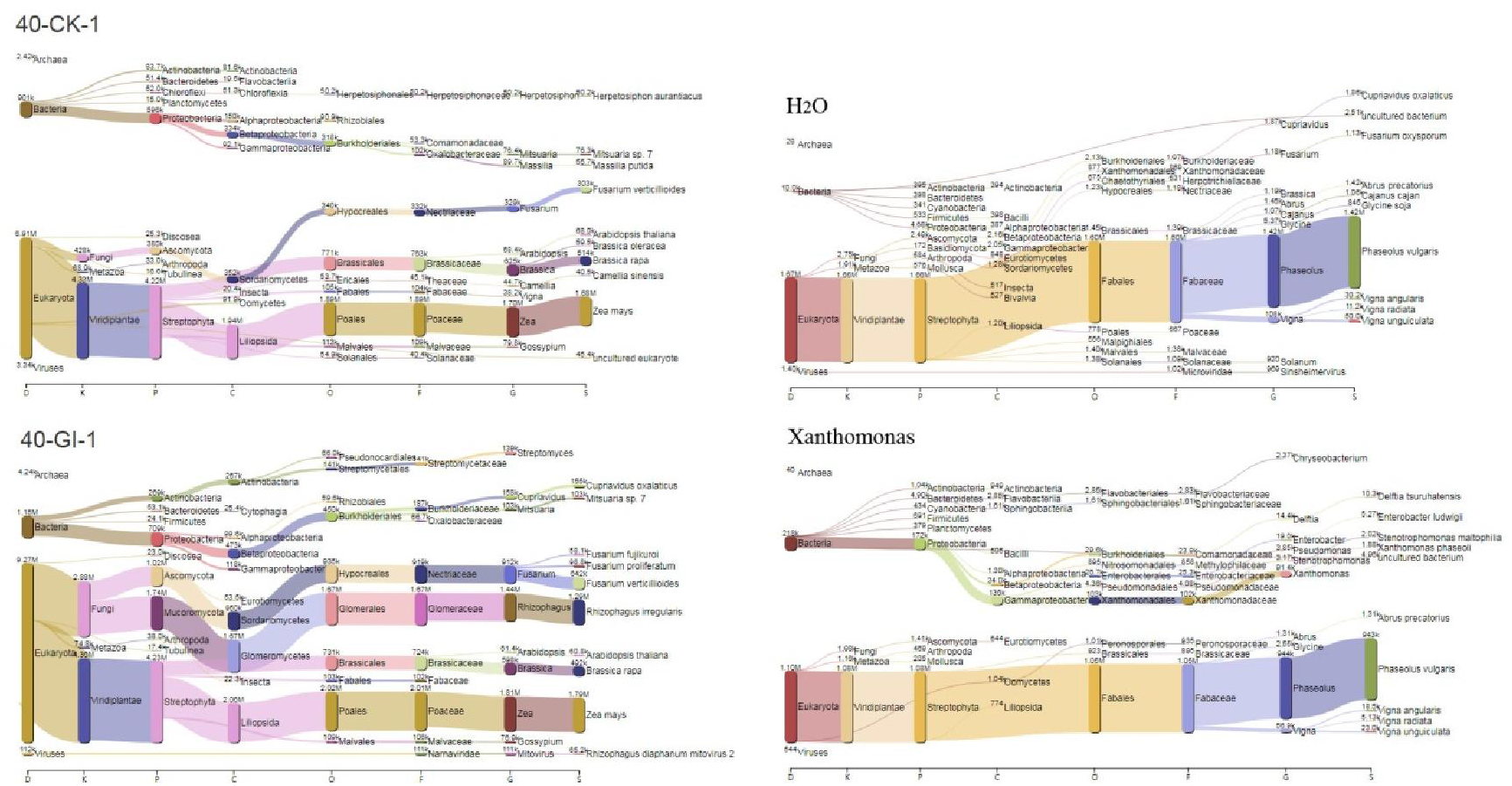
Species composition generated from unmapped files of the maize RNA-seq data and the common bean RNA-seq data, respectively. Note: 40-CK-1, one of the control samples of maize; 40-GI-1, one of the maize roots inoculated with *R. irregularis* DAOM-197198; H_2_O, one of the control samples of common bean; Xanthomonas, one of the common bean sample inoculated with *X. phaseoli* pv. *phaseoli* strain CFBP6546R

**Figure 4.**
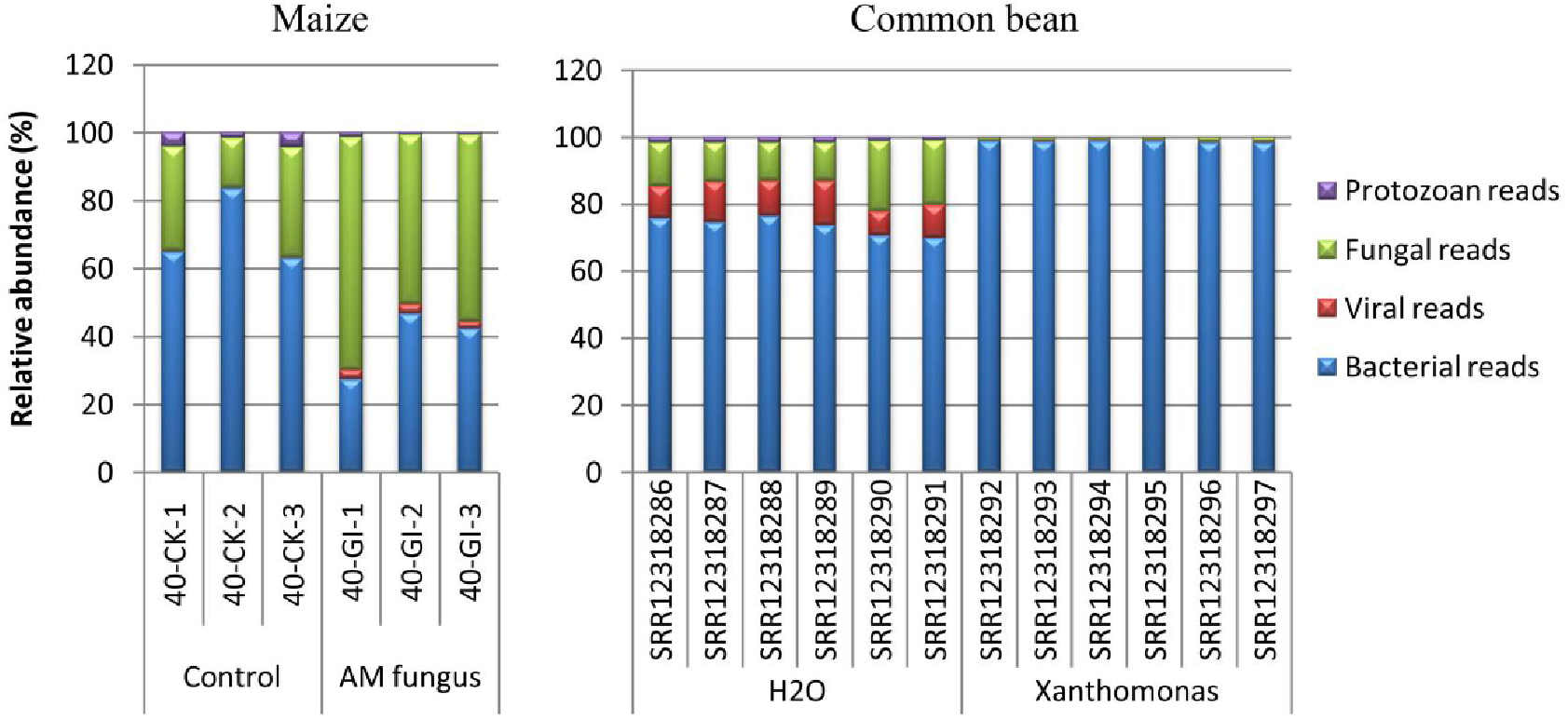
Relative abundance of different microorganisms generated from the maize RNA-seq data and the common bean RNA-seq data, respectively. Note: Ck, the control sample of maize; AM fungus, the maize roots inoculated with *R. irregularis* DAOM-197198; H2O, the control sample of common bean; Xanthomonas, the common bean sample inoculated with *X. phaseoli* pv. *phaseoli* strain CFBP6546R

In our previous study, the maize seedling was inoculated with *R. irregularis* DAOM-197198 to investigate the regulatory network responsive to AM fungi colonization in maize roots. The sankey diagram clearly shows that a large number of *R. irregularis* can be detected in the maize inoculated with AM fungus while the control was not detected (Figure 3). Similarly, *X. phaseoli* can be detected in the sample inoculated with *X. phaseoli* while the control was not detected (Figure 3). Species *R. irregularis* is the most significant biomarker in the maize roots inoculated with AM fungus *R. irregularis* (Figure 5), while Genus *Xanthomonas* is one of the most significant biomarkers in the common bean sample inoculated with *X. phaseoli* pv. *phaseoli* strain CFBP6546R (Figure 6). The evidence shows that the method of mining endophytic microbiome information from plant transcriptome data is reliable, and this method is also applicable to animal transcriptomes.

**Figure 5.**
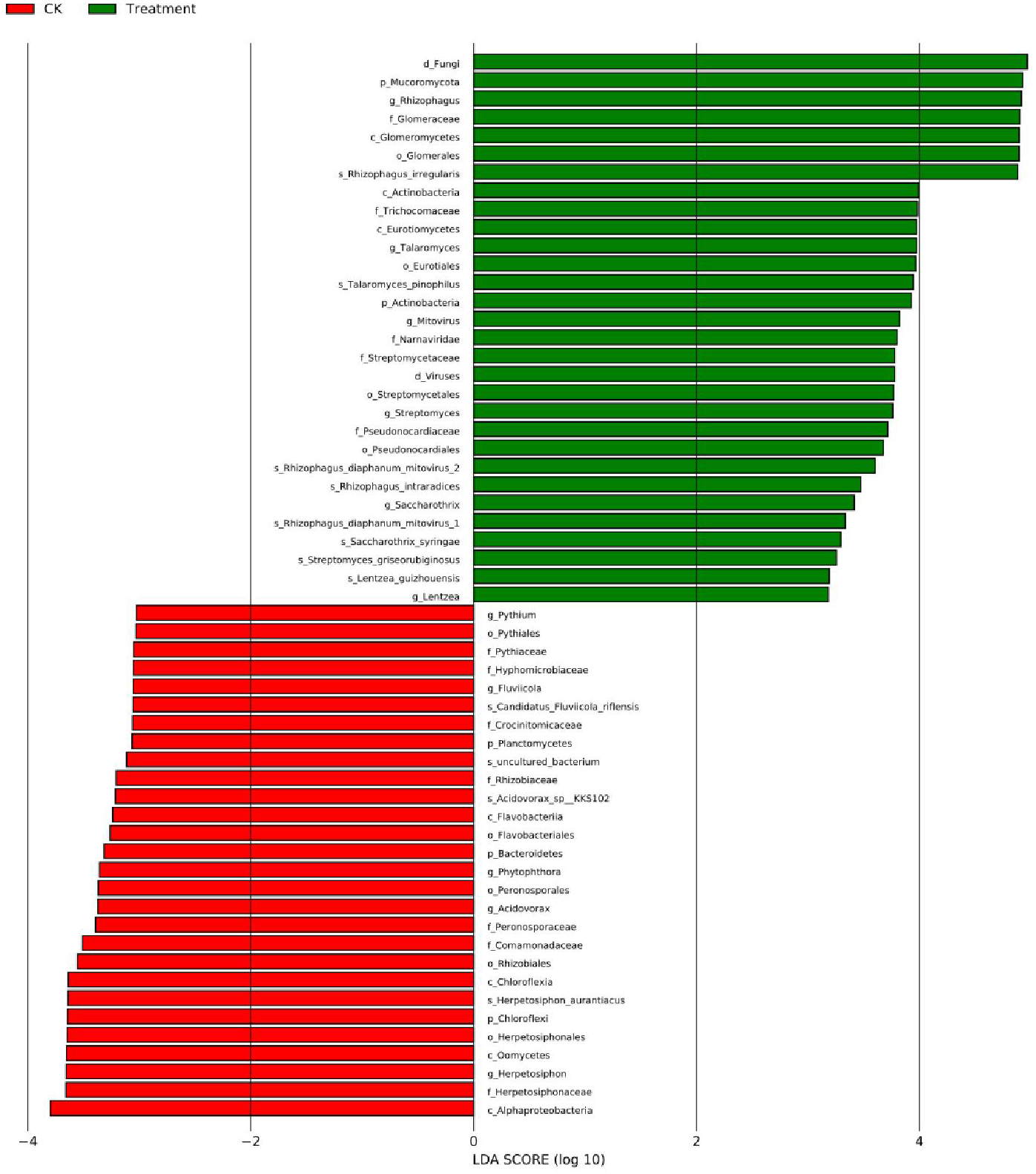
Taxons with significant difference between treatments in maize. Note: Ck, the control sample of maize; Treatment, the maize roots inoculated with *R. irregularis* DAOM-197198

**Figure 6.**
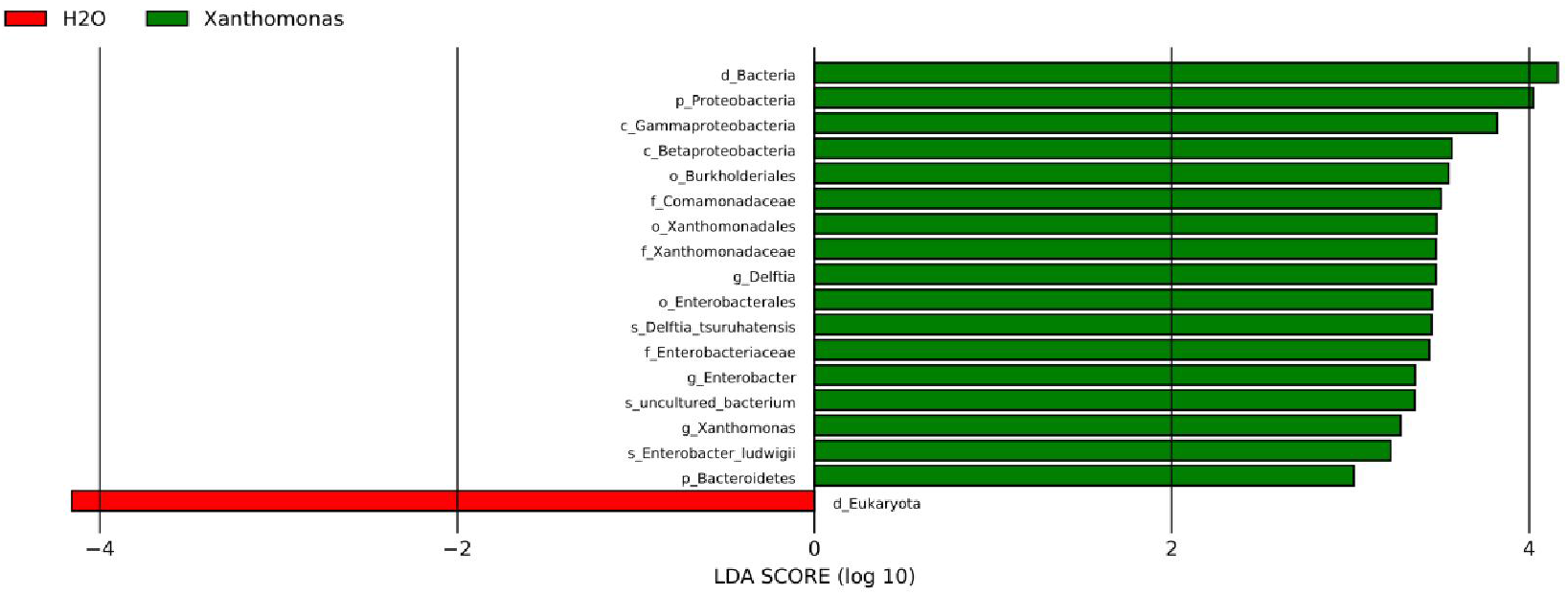
Taxons with significant difference between treatments in common bean. Note: H2O, the control sample of common bean; Xanthomonas, the common bean sample inoculated with *X. phaseoli* pv. *phaseoli* strain CFBP6546R

## Conclusions

We developed a method of mining endophytic microbiome information from plant or animal transcriptome data, which is reliable and useful in investigating microbiome information while investigating the gene expression information of the host plant or animal.

---

**Table S1.** The cleaning and mapping information of the maize RNA-seq data

| Treatment | Sample id | Input Read Pairs | Both Surviving Pairs | Aligned concordantly 0 times | Aligned concordantly exactly 1 time | Aligned concordantly >1 times |
| --- | --- | --- | --- | --- | --- | --- |
| CK | 40-CK-1 | 43293767 | 37896934 | 24.24% | 68.02% | 7.74% |
| CK | 40-CK-2 | 53402529 | 47289618 | 18.25% | 74.14% | 7.60% |
| CK | 40-CK-3 | 55730952 | 50485352 | 24.19% | 68.27% | 7.54% |
| AM fungus | 40-GI-1 | 51811515 | 45108797 | 28.26% | 64.73% | 7.00% |
| AM fungus | 40-GI-2 | 57804926 | 49615744 | 23.77% | 69.48% | 6.75% |
| AM fungus | 40-GI-3 | 53450977 | 46803974 | 23.27% | 69.84% | 6.89% |

Note: Ck, the control sample of maize; Treatment, the maize roots inoculated with *R. irregularis* DAOM-197198

**Table S2.** The cleaning and mapping information of the common bean RNA-seq data

| Treatment | Sample id | Input Read Pairs | Both Surviving reads | Aligned concordantly 0 times | Aligned concordantly exactly 1 time | >Aligned concordantly >1 times |
| --- | --- | --- | --- | --- | --- | --- |
| H2O | SRR12318286 | 29080436 | 28032538 | 5.65% | 90.78% | 3.57% |
| H2O | SRR12318287 | 30814752 | 29503109 | 5.94% | 90.53% | 3.53% |
| H2O | SRR12318288 | 30404194 | 29090030 | 5.27% | 90.76% | 3.96% |
| H2O | SRR12318289 | 31867409 | 30251534 | 5.61% | 90.47% | 3.91% |
| H2O | SRR12318290 | 26394779 | 25461066 | 6.51% | 90.68% | 2.80% |
| H2O | SRR12318291 | 27198114 | 26091099 | 6.81% | 90.39% | 2.80% |
| Xanthomonas | SRR12318292 | 15056487 | 14362639 | 10.14% | 86.96% | 2.89% |
| Xanthomonas | SRR12318293 | 14962446 | 14202256 | 10.72% | 86.45% | 2.83% |
| Xanthomonas | SRR12318294 | 18387872 | 17490043 | 11.04% | 85.69% | 3.27% |
| Xanthomonas | SRR12318295 | 19056924 | 18003970 | 11.64% | 85.17% | 3.19% |
| Xanthomonas | SRR12318296 | 16698533 | 16043346 | 9.28% | 86.48% | 4.24% |
| Xanthomonas | SRR12318297 | 16563339 | 15850911 | 9.86% | 85.96% | 4.18% |

Note: H2O, the control sample of common bean; Xanthomonas, the common bean sample inoculated with *X. phaseoli* pv. *phaseoli* strain CFBP6546R

## Notes

### Competing Interest Statement

The authors have declared no competing interest.

